# Long-term Exposure to Methyl Jasmonate Increases Myrosinases TGG1 and TGG2 in Arabidopsis *coi1* and *myc2,3,4* Mutants

**DOI:** 10.1101/2024.04.03.587911

**Authors:** Mohamadreza Mirzaei, Andisheh Poormassalehgoo, Kaichiro Endo, Shino Goto-Yamada, Ewa Dubas, Kenji Yamada

**Affiliations:** Malopolska Centre of Biotechnology, Jagiellonian University, Krakow, Poland; Doctoral School of Exact and Natural Sciences, Jagiellonian University, Krakow, Poland; The Franciszek Górski Institute of Plant Physiology, Polish Academy of Sciences, Krakow, Poland

**Keywords:** myrosinase, methyl jasmonate, COI1, myrosin cell

## Abstract

Plants are constantly subjected to stresses such as wounding or herbivore attacks, which continuously activate the JA signaling pathway in nature. However, knowledge about the effect of long-term activation of the JA signaling pathway on plant defense response remains limited. THIOGLUCOSIDE GLUCOHYDROLASE 1 (TGG1) and TGG2 are enzymes that activate defensive metabolites, namely glucosinolates, which are involved in defense against herbivores and pathogens. Here, we show that prolonged exposure to the wounding hormone methyl jasmonate (MeJA) enhances *TGG1* and *TGG2* expression independent of the canonical jasmonic acid (JA) signaling pathway in *Arabidopsis thaliana* rosette leaves. We found that airborne MeJA treatment for up to 5 d enhanced both *TGG1* and *TGG2* gene expression and their protein levels in Arabidopsis leaves. Notably, *TGG1* and *TGG2* gene expressions significantly upregulated in two JA signaling pathway mutants, namely *coi1-16* and *myc2,3,4*, following 5 d of MeJA treatment. TGG1 and TGG2 proteins accumulate in specialized myrosin cells of rosette leaves. Myrosin cell area expands in response to MeJA treatment in a leaf-age-dependent manner. Consistent with this, the expression of *FAMA*, a transcription factor known to regulate *TGG1* and *TGG2* gene expression, also increased in a leaf-age-dependent manner after 5 d of MeJA treatment. Taken together, our results suggest the existence of a non-canonical JA signaling pathway that is activated by long-term exposure to MeJA and regulates the expression of defense-related genes *TGG1* and *TGG2*.

## 1 Introduction

The myrosinase-glucosinolate system is a chemical defense mechanism against herbivores and pathogens in plants of the order Brassicales. This defense mechanism was first discovered in mustard seeds (Bussy, 1840; Bones and Rossiter, 1996) and consists of an enzyme called myrosinase and a family of substrates called glucosinolate. Myrosinases and glucosinolates are stored in different tissues or subcellular compartments in plants (Shirakawa et al., 2016). However, these enzymes and substrates come into contact when tissue damage occurs, and the enzymes begin to hydrolyze glucosinolates to produce toxic compounds, such as isothiocyanateto stop further damage by herbivores and pathogens (Bhat & Vyas, 2019). In *Arabidopsis thaliana*, canonical myrosinase activity is encoded by six *THIOGLUCOSIDE GLUCOHYDROLASE* (*TGG*) genes, namely *TGG1*–*TGG6* (Rask et al., 2000; Shirakawa & Hara-Nishimura, 2018). Two of the six genes, *TGG1* (*At5g26000*) and *TGG2* (*At5g25980*), are expressed in the leaves, whereas *TGG4* (*At1g47600*) and *TGG5* (*At1g51470*) are expressed in the roots (Shirakawa et al., 2016; Andersson et al., 2009). *TGG3* (*At5g48375*) and *TGG6* (*At1g51490*) were considered pseudogenes because of their open reading frame disruption (Wang et al., 2009).

In *Arabidopsis*, TGG1 and TGG2 proteins are stored in two types of cells with similar ‘idioblast’ characteristics: myrosin cells and stomata guard cells (Shirakawa et al., 2022). Myrosin cells are located next to phloem cells, and this particular name was given because myrosinases accumulate extensively in their vacuoles (Guignard, 1980; Heinricher, 1884; Bones and Rossiter, 1996). Although myrosin cells are localized adjacent to the phloem, they develop independently of vascular precursor cells and originate directly from ground meristem cells (Shirakawa et al., 2014; Shirakawa et al., 2016). Basic helix-loop-helix (bHLH) transcription factors (TFs) FAMA, SCREAM1 and SCREAM2, which regulate stomatal guard cell differentiation, regulate the differentiation of myrosin cells (Li and Sack, 2014; Shirakawa et al., 2014). Although myrosinases have a chemical defense function in both myrosin and stomatal cells, it is reported that glucosinolate metabolism mediated by TGG1 in the stomatal cells plays a role in stomatal movement (Bones and Rossiter, 1996; Zhang et al., 2019).

Jasmonic acid (JA) and its metabolic derivatives, such as jasmonoyl-isoleucine (JA-Ile) and methyl jasmonate (MeJA), are cyclopentanone derivatives of linolenic acid (Devoto et al., 2002; Liu & Timko, 2021). These chemicals are widely distributed phytohormones in higher plants and play important roles in the stress response and regulation of several plant developmental processes (Ruan et al., 2019). In the JA signaling pathway, JA-Ile is sensed by CORONATINE INSENSITIVE 1 (COI1)-JASMONATE ZIM-DOMAIN (JAZ) co-receptors (Xie et al., 1998; Devoto et al., 2002). COI1, a leucine-rich repeat (LRR)-F-box protein, forms the Skp1–Cul1–F-box-protein (SCF) ubiquitin ligase complex, which is called SCF^COI1^. The hormone ligand promotes binding between SCF^COI1^ and JAZ proteins for the ubiquitination of JAZ proteins and their subsequent degradation by the 26S proteasome. In the non-JA-signaling state, JAZ proteins physically interact with MYC2/3/4 TFs and repress their functions (Chini et al., 2009; Ruan et al., 2019). JA-induced JAZ protein degradation releases downstream TFs that activate various JA-inducible genes. In *Arabidopsis*, JA-inducible genes include *VEGETATIVE STORAGE PROTEIN 2* (*VSP2*), *JAZ2, JASMONATE RESPONSIVE 1* (*JR1*), and *BETA GLUCOSIDASE 18* (*BGLU18*), which are used for the analysis of JA response (Liu & Timko, 2021; Schweizer et al., 2013; Stefanik et al., 2020).

It has been reported that myrosinase activity is significantly lower in the *coi1* mutant than in the wild type when sinigrin is used as a substrate (Capella et al., 2001), and MeJA treatment increases *TGG1* expression in a COI1-dependent manner in *Arabidopsis* (Feng et al., 2021). Plants frequently encounter stress such as wounding or herbivore attacks, which continuously induce JA biosynthesis and activate the JA signaling pathway (McConn et al., 1997; Halitschke and Baldwin, 2004). Because continuous herbivore attacks are usual in natural conditions, the analysis of short-term MeJA response in the laboratory experiment may not explain the entire plant response. Therefore, to better understand the plant response under continuous stress, the present study investigated the JA responses of *TGG1* and *TGG2* under continuous long-term MeJA treatment in wild-type and mutant *Arabidopsis* lacking key JA signaling pathway components, namely, *coi1-16* and *myc2,3,4*. This approach allows for a deeper understanding of how these genes function under conditions more like those found in nature. We unexpectedly found that although *TGG1* and *TGG2* are JA-responsive genes, the response is not fully dependent on canonical JA signaling pathway components, such as COI1 and MYC2/3/4, when plants are continuously exposed to MeJA for a long time.

## 2 Materials and Methods

### 2.1 Plant materials, plant growth condition

The *Arabidopsis thaliana* Col-0 accession was used as the wild-type. Transgenic plants containing *ProTGG2:VENUS-2sc* and *pFAMA:GFP* constructs were kind gifts from M. Shirakawa and I. Hara-Nishimura (Shirakawa et al., 2014; Shirakawa and Hara-Nishimura, 2018). The *myc2,3,4* mutant was kindly provided by H. Frerigmann, and the *coi1-16* mutant was purchased from the *Arabidopsis* Biological Resource Center (ABRC). The *myc2,3,4* mutant was generated by crossing T-DNA insertion mutants *myc4* (GK491E10), *myc3* (GK445B11), and *myc2*/*jin1-9* (SALK_017005) (Frerigmann et al., 2014). The *coi1-16* mutant has a point mutation in the *COI1* gene, resulting in an amino acid alteration of L245F (Ellis and Turner, 2002). All mutants were of the Columbia ecotype. Following sterilization with 70% (v/v) ethanol, seeds were cultivated at 4 cm from the center of the plate (Supplementary Fig. 1) and incubated at 4 °C for 2 d prior to germination. Seeds were then germinated at 22 °C under continuous light (approximately 100 µE s^-1^ m^-2^) in half strength Murashige and Skoog (1/2 MS) medium [1/2 MS basal salt mixture (092623020, MP Biomedicals), 1% (w/v) sucrose and 0.5% (w/v) MES-KOH (pH 5.7)] containing 0.4% (w/v) Gellan Gum (Wako, Japan). Leaves were numbered according to ontogenetic development, starting with the first true leaf, and analyzed (Fig. 1h).

**Figure 1.**
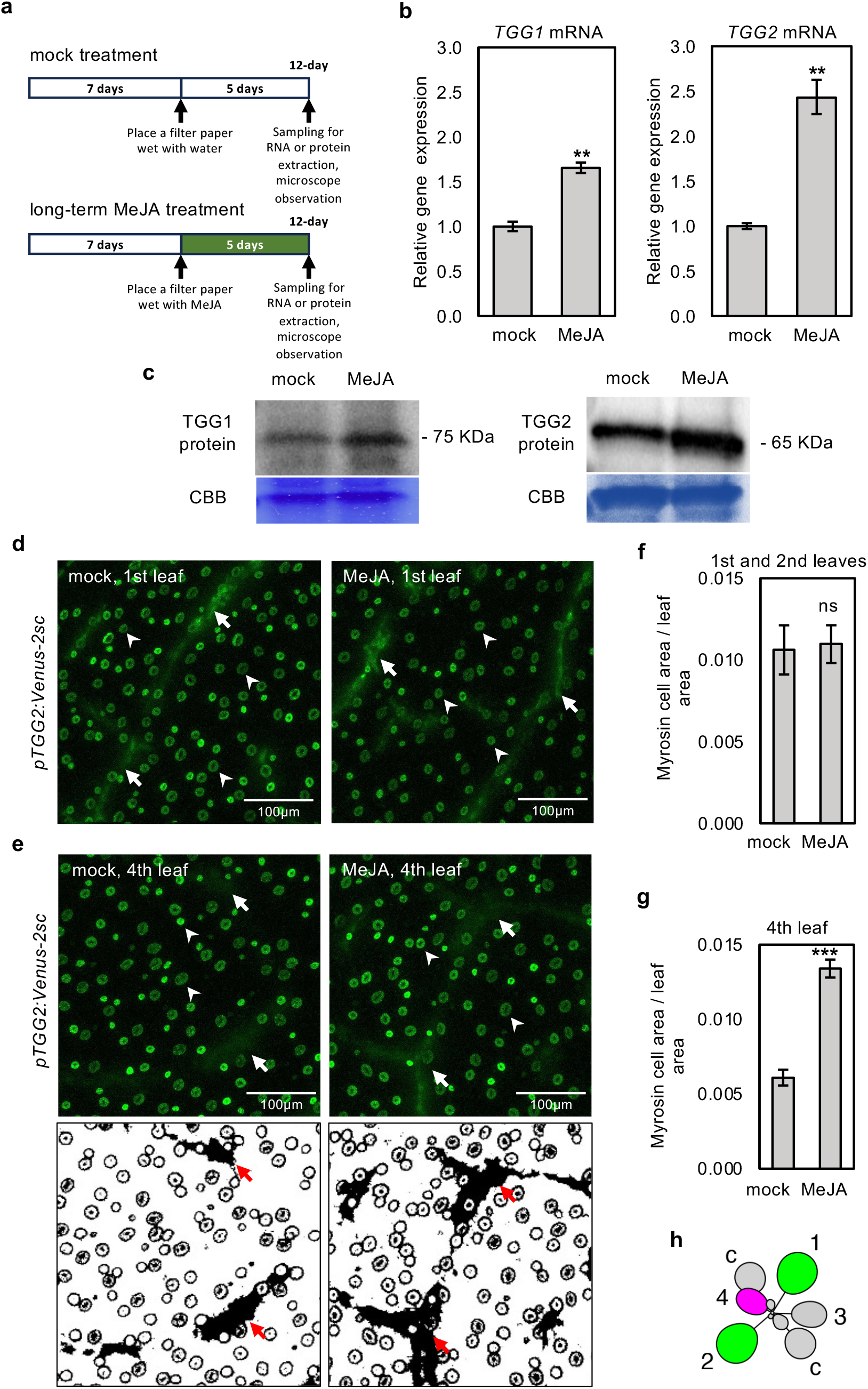
Long-term MeJA treatment increases the expression of *TGG1* and *TGG2* in wild-type rosette leaves. **a)** The schemes show the experimental setup of mock and MeJA treatment. MeJA treatment was started 7 d after germination, and plants were sampled 12 d after germination. **b, c)** The relative expression levels of *TGG1* and *TGG2* (b), and the protein amount of TGG1 and TGG2 (c) in the shoots of mock or long-term MeJA treated *Arabidopsis* wild type (Col-0) plants. Error bars denote the standard error of three biological replications. Double asterisks denote *p* < 0.01 based on the student’s *t*-test. **d, e)** Confocal microscopic images of the first (d) and fourth (e) leaves of 12-d-old transgenic plants harboring *pTGG2:Venus-2sc*. Myrosin cells (arrows) and stomata guard cells (arrowheads) are recognized with Venus fluorescence but have different shapes. The processed images show myrosin cells as a black area (e, lower, red arrows). **f, g)** The ratio of myrosin cell areas per leaf area in the first and second (f) or fourth (g) leaves, which is calculated from the images (e, lower). Error bars denote the standard error of seven biological replications. *** denotes *p* < 0.001 and ns denotes no significance based on the student’s *t*-test. **h)** The leaf numbering. c, cotyledons.

### 2.2 MeJA treatment

The liquid of MeJA (500 nmol/plate) (Sigma-Aldrich) was applied to a filter paper pad left in the cap of an Eppendorf tube (1.5 mL) for evaporation and positioned in the center of a plate (φ = 9 cm) (Supplementary Fig. 1). The evaporated MeJA gradually and uniformly diffused into the enclosed air space of the plate with a saturation concentration of around 1.6 µg/L (Acevedo et al, 2003). The distance between the plants and the source of MeJA was important, and the arrangement ensured uniform and equal exposure of all seedlings to MeJA. Leaf samples for each replicate were randomly collected, frozen in liquid nitrogen, and stored at –80 °C for subsequent molecular analyses.

### 2.3 Binocular and confocal microscope

A binocular fluorescence microscope (Zeiss SteREO V12) was used to observe fluorescent proteins in whole leaves. The images were captured using a CCD camera (Zeiss AxioCam MRc5). A confocal laser scanning microscope (LSM880, Carl Zeiss, Jena, Germany) was used to observe fluorescent proteins.

### 2.4 RNA isolation and Quantitative RT-PCR

Total RNA was extracted from the aerial parts (aboveground) of 12-d-old seedlings using the TRI Reagent (Molecular Research Center, Cincinnati, USA). After dissolving the RNA in distilled water and digesting contaminated genomic DNA with DNase I (Sigma-Aldrich), the first cDNA strand was generated from 1 µg of the RNA using Ready-to-Go RT-PCR beads (GE Healthcare) and random oligomers. PowerUp SYBR Green Master Mix (Thermo Fisher Scientific, USA) was used for the quantification of cDNA of *UBIQUITIN 10* (*UBQ10*), *TGG1, TGG2, FAMA, VSP2, BGLU18, JR1*, and *JAZ2* using an instrument (QuantStudio 12K Flex, Thermo Fisher Scientific). Gene-specific primer sets were generated using the Primer3Plus software (Supplementary Table 1). The *UBQ10* gene was used as a reference gene because of its stable and constitutive expression in all *Arabidopsis* tissues and treatments (Czechowski et al., 2005). Relative expression of the target genes was normalized to that of *UBQ10*.

### 2.5 SDS-PAGE and Immunoblot Analysis

Total proteins were extracted from 50 mg of leaves with 200 µL of 2× sample buffer (20 mM Tris-HCl buffer, pH 6.8, 40% (v/v) glycerol, 2% (w/v) sodium dodecyl sulfate (SDS), and 2% (v/v) 2-mercaptoethanol). The homogenate was centrifuged at 12,000 × *g* for 5 min to remove debris. Extracts (10 µL) were electrophoresed on SDS-polyacrylamide gels. After separation, the proteins were transferred onto a nylon membrane and subjected to immunoblotting. The anti-TGG1 and anti-TGG2 antibodies were diluted 5000-fold for each treatment (Ueda et al., 2006; Shirakawa et al., 2014). The proteins were stained with Coomassie Brilliant Blue R-250.

### 2.6 Insect feeding assays

Tropical house crickets (*Gryllodes sigillatus*) of the same age, originating from the same colony, were purchased from the market. The crickets were starved for 2 d before the experiments. The 7-d-old *Arabidopsis* wild type plants, *coi1-16* and *myc2,3,4* mutants were treated with either mock or MeJA for 5 d. Afterward, ten MeJA-treated and ten untreated plants were carefully removed from their growing medium and placed in a 20×20×20 cm box for the dual-choice feeding assay. To prevent desiccation of plants, their roots were wrapped in wet tissue paper. Ten starved crickets were released into each box and allowed to consume the plants by keeping them overnight. Plant weights were recorded before and after the feeding experiments.

### 2.7 Accession Numbers

Sequence data from this study can be obtained in the *Arabidopsis* Genome Initiative databases under the following accession numbers: *TGG1* (*At5g26000*), *TGG2* (*At5g25980*), *FAMA* (*At3g24140*), *COI1* (*At2g39940*), *MYC2* (*At1g32640*), *MYC3* (*At5g46760*), *MYC4* (*At4g17880*), *VSP2* (*At5g24770*), *BGLU18* (*At1g52400*), *JR1* (*At3g16470*), *JAZ2* (*At1g74950*), and *UBQ10* (*At4g05320*).

## 3 Results

### 3.1 Long-term exposure to airborne MeJA increases the expression level of *TGG1* and*TGG2* in *Arabidopsis* rosette leaves

JA treatment inhibits true leaf and cotyledon growth (Zhang and Turner, 2008), elongation (Chen et al., 2013), and adventitious root development (Gutierrez et al., 2012). We treated 7-d-old *Arabidopsis thaliana* seedlings with vaporized MeJA for 5 d (Fig. 1a) and found that airborne MeJA treatment reduced plant size (Supplementary Fig. 1), as previously reported (Zhang and Turner, 2008; Chen et al. 2013; Gutierrez et al., 2012). Next, we examined the expression levels of two myrosinase genes, *TGG1* and *TGG2*, after airborne MeJA treatment to determine whether JA induces the expression of these genes. Reverse transcription-quantitative PCR (RT-qPCR) analysis of shoots revealed that the expression levels of both genes were significantly increased after 5 d of airborne MeJA treatment (Fig. 1b). In addition, immunoblot analysis showed that exposure to airborne MeJA for 5 d significantly increased TGG1 and TGG2 protein levels in the shoots (Fig. 1c).

To monitor the activity of the *TGG2* promoter in each rosette leaf, we used a transgenic reporter line containing the *TGG2* promoter and *Venus-2sc* reporter construct (*pTGG2:Venus-2sc*), in which Venus fluorescence was observed mainly in myrosin cells and to some extent in the stomata guard cells (Fig. 1d) (Shirakawa et al., 2016). We measured the myrosin cell area in the images based on Venus fluorescence. We divided the leaves into two types based on the leaf developmental stage at the starting point of MeJA treatment: 1) old (first and second) leaves that have completed leaf development, and 2) young (fourth) leaves that are in the course of leaf development. The myrosin cell area was not dramatically changed by MeJA treatment in the first and second leaves, but was significantly increased in the fourth leaves of MeJA-treated plants (Fig. 1e, f, g). The Venus fluorescence intensity in myrosin cells did not change significantly (Supplementary Fig. 2). We also measured the myrosin cell area of the eleventh leaves after very long-term MeJA treatment. The myrosin cell area of these young leaves tended to increase after a very long MeJA treatment (Supplementary Fig. 3). These findings suggest that MeJA promoted myrosin cell area expansion in young leaves at certain plant ages or leaf orders. The increase in myrosin cell area can be attributed to either an increase in the number of myrosin cells or an increase in the volume of myrosin cells. Because we could not count myrosin cells in the fourth leaves, we used the emerging first leaves of younger seedlings to observe the changes in myrosin cell number after MeJA treatment. We started MeJA treatment on an earlier day (2 d after germination) when the first leaf was emerging. Microscopic observations demonstrated that the myrosin cell number was not increased by MeJA treatment in the emerging first leaf (Supplementary Fig. 4). These findings suggest that MeJA treatment increases myrosin cell area by presumably increasing each myrosin cell volume but not by promoting myrosin cell proliferation.

As TGG2 is known to accumulate in stomatal guard cells, we counted the number of stomatal cells in MeJA-treated leaves. No significant changes in stomatal density were observed in the first, second, and fourth leaves (Supplementary Fig. 5).

### 3.2 Long-term airborne MeJA treatment increases *TGG1* and *TGG2* expression in the canonical JA-insensitive *coi1*-*16* and *myc2,3,4* mutants

COI1 is the JA receptor that forms a co-receptor complex with JAZ proteins, and MYC2, MYC3, and MYC4 are key regulators that control the expression of JA response genes (Sheard et al., 2010; Fernandez-Calvo et al., 2011). To examine whether the canonical JA-signaling pathway components regulate the expression of *TGG1* and *TGG2*, we examined their expression levels in *Arabidopsis coi1* and *myc2,3,4* deficient mutants. We used the *coi1-16* mutant allele because it is fertile at 16 °C, unlike other *coi1* mutant alleles, and can be maintained as a pure homozygous line (Ellis and Turner, 2002). Long-term (5 d) airborne MeJA treatment led to a significant increase in *TGG1* and *TGG2* expression levels in the wild-type plants. Interestingly, this response was also observed in *coi1-16* and *myc2,3,4* mutants (Fig. 2). We examined TGG1 and TGG2 protein levels by immunoblot analysis, and the results showed that short-term (1 d) MeJA treatment did not increase TGG1 and TGG2 accumulation, but long-term treatment increased these protein levels in wild type, *coi1-16* and *myc2,3,4* plants, which was consistent with *TGG1* and *TGG2* transcript levels (Fig. 3). Taken together, our results suggest that long-term MeJA treatment induces *TGG* genes independently of the canonical JA signaling pathway, including COI1 and MYC2/3/4.

**Figure 2.**
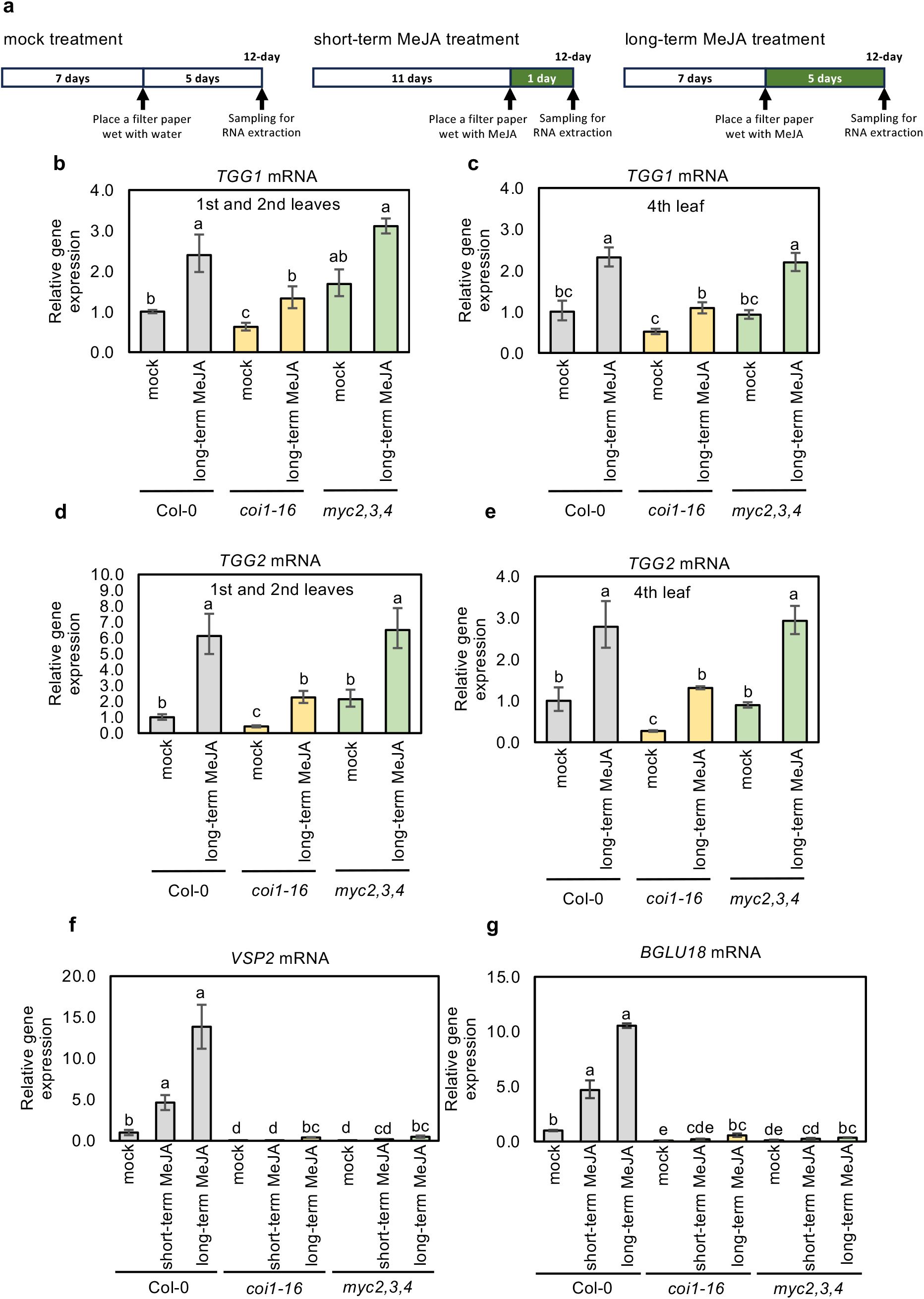
Long-term MeJA treatment increased the expression of *TGG1* and *TGG2* in the first, second, and fourth leaves of Col-0, *coi1-16* and *myc2,3,4* mutants, while the expression of canonical MeJA-responsive genes decreased significantly in *coi1-16* and *myc2,3,4* mutants. **a)** The schemes show the experimental setup of mock, short-term and long-term MeJA treatments. MeJA treatment was started 7 (for the long-term MeJA treatment) or 11 d (for the short-term MeJA treatment) after germination, and plants were sampled 12 d after germination. **b-e)** The relative expression levels of *TGG1* (b, c) and *TGG2* (d, e) in the first and second (b, d), and fourth (c, e) leaves of mock and long-term MeJA treated Arabidopsis wild type (Col-0), *coi1-16* and *myc2,3,4* mutants. **f,g)** The relative expression levels of *VSP2* (f) and *BGLU18* (g) in the shoots of mock, short- and long-term MeJA-treated Arabidopsis wild type (Col-0), *coi1-16* and *myc2,3,4* mutants. Error bars denote the standard error of three biological replications. Different lowercase letters indicate significant differences (p < 0.05; Tukey’s test).

**Figure 3.**
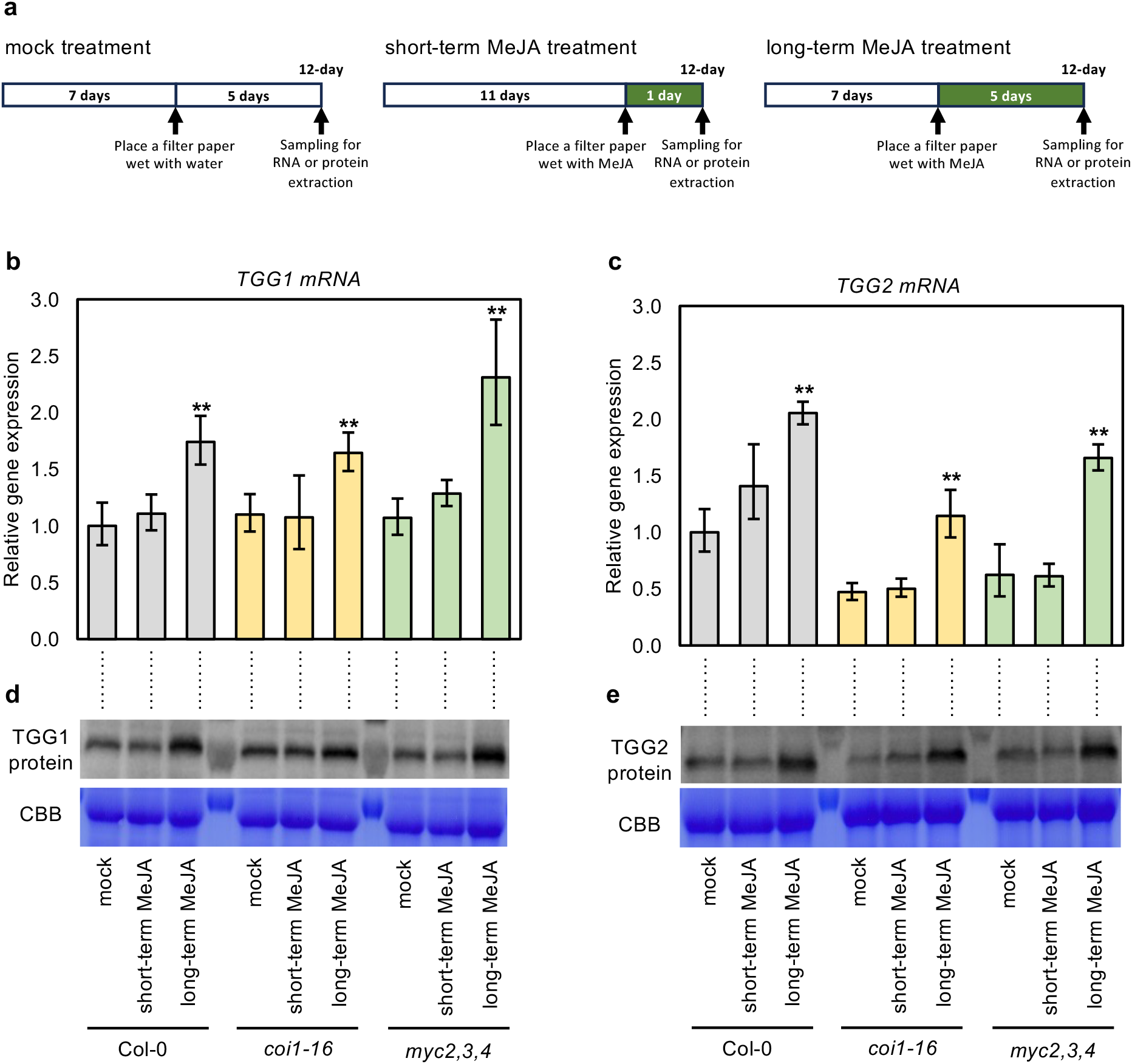
Long-term MeJA treatment increases the expression of TGG1 and TGG2 in *coi1-16* and *myc2,3,4* mutants. **a)** The schemes show the experimental setup of mock, short-term and long-term MeJA treatments. MeJA treatment was started 7 (for the long-term MeJA treatment) or 11 d (for the short-term MeJA treatment) after germination, and plants were sampled 12 d after germination. **b-e)** The relative expression levels of *TGG1* (b) and *TGG2* (c), and the protein amount of TGG1 (d) and TGG2 (e) in the shoots of mock, short- and long-term MeJA treated Arabidopsis wild type (Col-0), *coi1-16* and *myc2,3,4* mutants. Error bars denote the standard error of three biological replications. Double asterisks denote *p* < 0.01 based on the student’s *t*-test.

Recently, Feng et al. (2021) reported that 6 h MeJA treatment upregulates the *TGG1* expression in a COI1-dependent manner. To examine how long treatment is required for the stimulation of COI1-independent *TGG1* and *TGG2* expression with airborne MeJA treatment, we conducted a time-course experiment by changing the starting date for the MeJA treatment in wild-type and *coi1-16* plants. The result revealed that both *TGG1* and *TGG2* expressions were significantly increased after 2 d MeJA treatment in wild type, and after 4 d for *TGG1* and 2 d for *TGG2* in *coi1-16* mutant (supplementary Fig. 6).

As the *coi1-16* and *myc2,3,4* mutants have been described to have a leaky JA response (Ellis and Turner, 2002; Song et al., 2017), and the extent of the JA response in these mutants was not known in the long-term airborne MeJA treatment, we examined the plant growth phenotype of these mutants in the long-term airborne MeJA treatment as the same experimental condition as in Fig. 1a. Airborne MeJA treatment reduced shoot and root growth and shoot fresh weight (Supplementary Fig. 7) in the wild type, but not in the *coi1-16* and *myc2,3,4* mutants. These results indicate that *coi1-16* and *myc2,3,4* mutants reduced or eliminated the MeJA response under our experimental conditions.

To further investigate the response of *coi1-16* and *myc2,3,4* mutants to airborne MeJA treatment, we examined the expression levels of the known COI1-dependend JA-inducible genes *VSP2, BGLU18, JR1*, and *JAZ2* (Boter et al., 2004; Chung et al., 2009). The expression levels of *VSP2* and *BGLU18* were significantly increased after short-term (1 d) airborne MeJA treatment, and these levels were further increased by long-term (5 d) airborne MeJA treatment in wild-type plants. In contrast, *coi1-16* and *myc2,3,4* mutant seedlings showed strongly reduced expression of *VSP2* and *BGLU18* under the same treatment (Fig. 2f, g). Similar results were observed for *JR1* and *JAZ2*, and their expressions were significantly lower in *coi1-16* and *myc2,3,4* mutants than in wild-type plants after long-term MeJA treatment (Supplementary Fig. 8). These results indicate that *VSP2, BGLU18, JR1*, and *JAZ2* expressions are dependent on COI1 and MYC2/3/4, suggesting that the pathway regulating *TGG1* and *TGG2* expression is distinct from that regulating the *VSP2, BGLU18, JR1*, and *JAZ2* under long-term MeJA treatment.

### 3.3 Long-term MeJA treatment increased *FAMA* expression

Because FAMA is a regulator of myrosin cell differentiation and *TGGs* expression (Li and Sack, 2014; Shirakawa et al., 2014; Feng et al., 2021), we further investigated the possible role of MeJA in increasing myrosin cell differentiation by monitoring the expression of *FAMA*. Long-term MeJA treatment upregulated the expression of *FAMA* in the first, second, and fourth leaves of both wild-type and *coi1-16* mutant plants, while no significant change in *FAMA* expression was observed in *myc2,3,4* mutant (Fig. 4a, b; Supplementary Fig. 9). *FAMA* is expressed in both stomatal guard and myrosin cells. To examine the *FAMA* expression patterns in stomatal guard cells and myrosin cells, we used *pFAMA:GFP* transgenic plants (Shirakawa and Hara-Nishimura, 2018). Long-term MeJA treatment increased the GFP fluorescence intensity of stomatal guard cells in the first, second, and fourth leaves, indicating an increase in *FAMA* expression in these cells (Supplementary Fig. 10). Regarding the myrosin cells area, a significant increase was observed in the fourth leaves of 12-d-old MeJA-treated plants compared to mock plants, but no *FAMA* expression was observed in the myrosin cells of the first and second leaves regardless of MeJA treatment (Fig. 4c-f). These findings suggest that the increase in *FAMA* expression in the first and second leaves of MeJA-treated plants is attributable to its increased expression in stomatal cells, whereas the increase in *FAMA* expression in the fourth leaf is a consequence of its higher expression in stomatal and myrosin cells.

**Figure 4.**
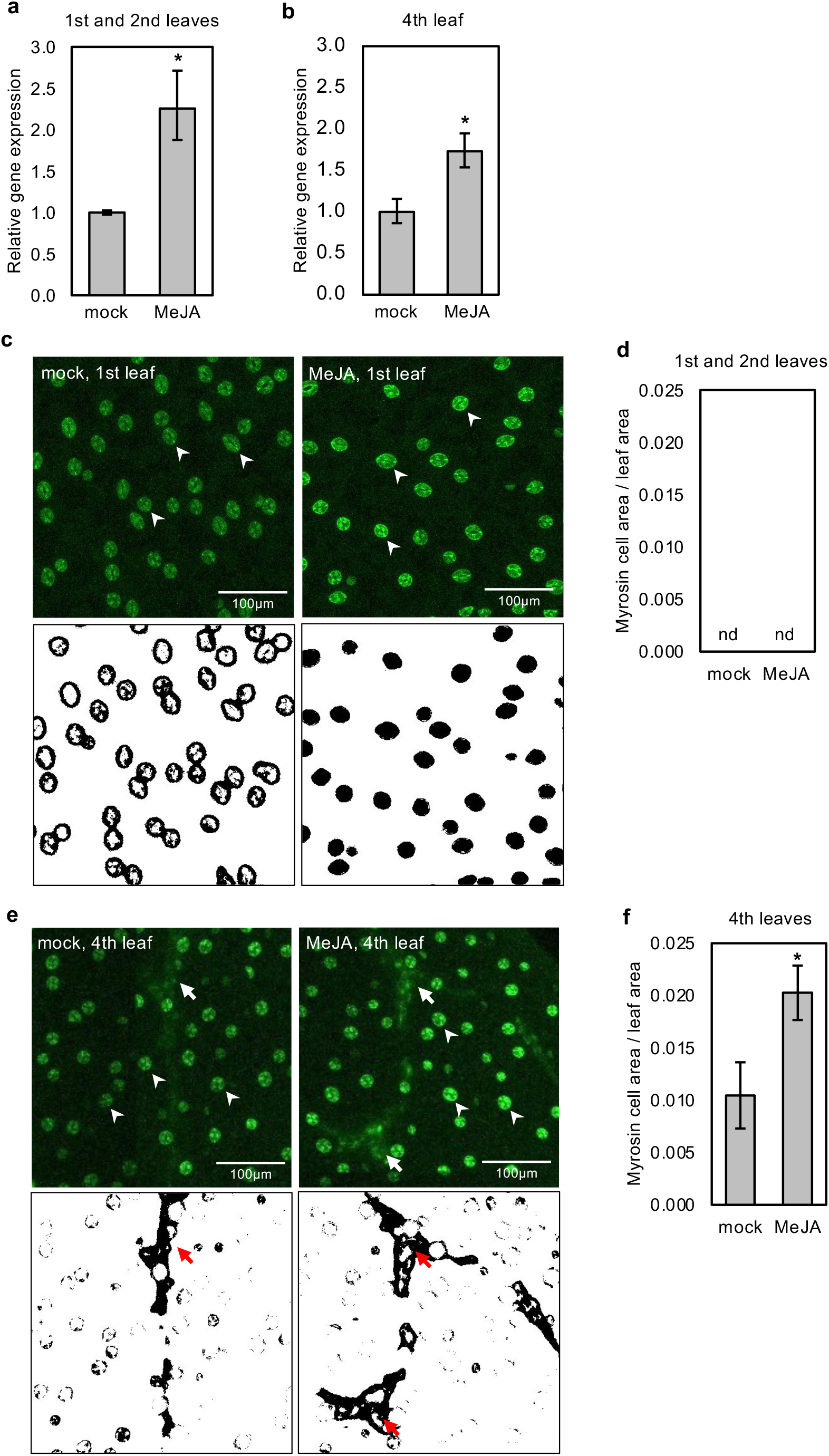
Long-term MeJA treatment increased the expression of *FAMA* in the first, second, and fourth leaves. **a, b)** The relative expression levels of *FAMA* in different rosette leaves of 12-d-old *Arabidopsis* wild-type (Col-0) plants. Error bars denote the standard error of three biological replications. * denotes *p* < 0.05 based on the student’s *t*-test. **c, e)** Confocal microscopic images (upper) of the first (c) and fourth (e) leaves of 12-d-old transgenic plants harboring *pFAMA:GFP*. Arrows and arrowheads represent myrosin cells and stomata guard cells, respectively. The processed images (lower) show myrosin cells as a black area (red arrows). **d, f)** The ratio of GFP-expressing myrosin cell areas per leaf area in the first and second (d) or fourth (f) leaves, which is calculated from the images (c, e, lower). Error bars denote the standard error of five biological replications. * denotes *p* < 0.05 based on the student’s *t*-test.

### 3.4 Long-term MeJA treatment decreased the feeding preference of crickets

To examine the effect of long-term MeJA treatment on defense against herbivores, we performed dual-choice feeding assays using crickets (*Gryllodes sigillatus*). The 7-d-old wild-type plants and *coi1-16* and *myc2,3,4* mutants were either mock- or MeJA-treated for 5 d before being exposed to starved crickets. In wild-type plants, long-term MeJA treatment reduced cricket feeding compared to that of mock-treated plants (Fig. 5 and Supplementary Fig. 11), indicating that long-term MeJA treatment enhanced plant defense against herbivores. In addition, surprisingly, the *coi1-16* significantly increased defense against herbivores following long-term MeJA treatment, and this trend was also observed in the *myc2,3,4* mutant (Fig. 5 and Supplementary Fig. 11), suggesting that long-term MeJA treatment enhanced herbivory resistance in these mutants.

**Figure 5.**
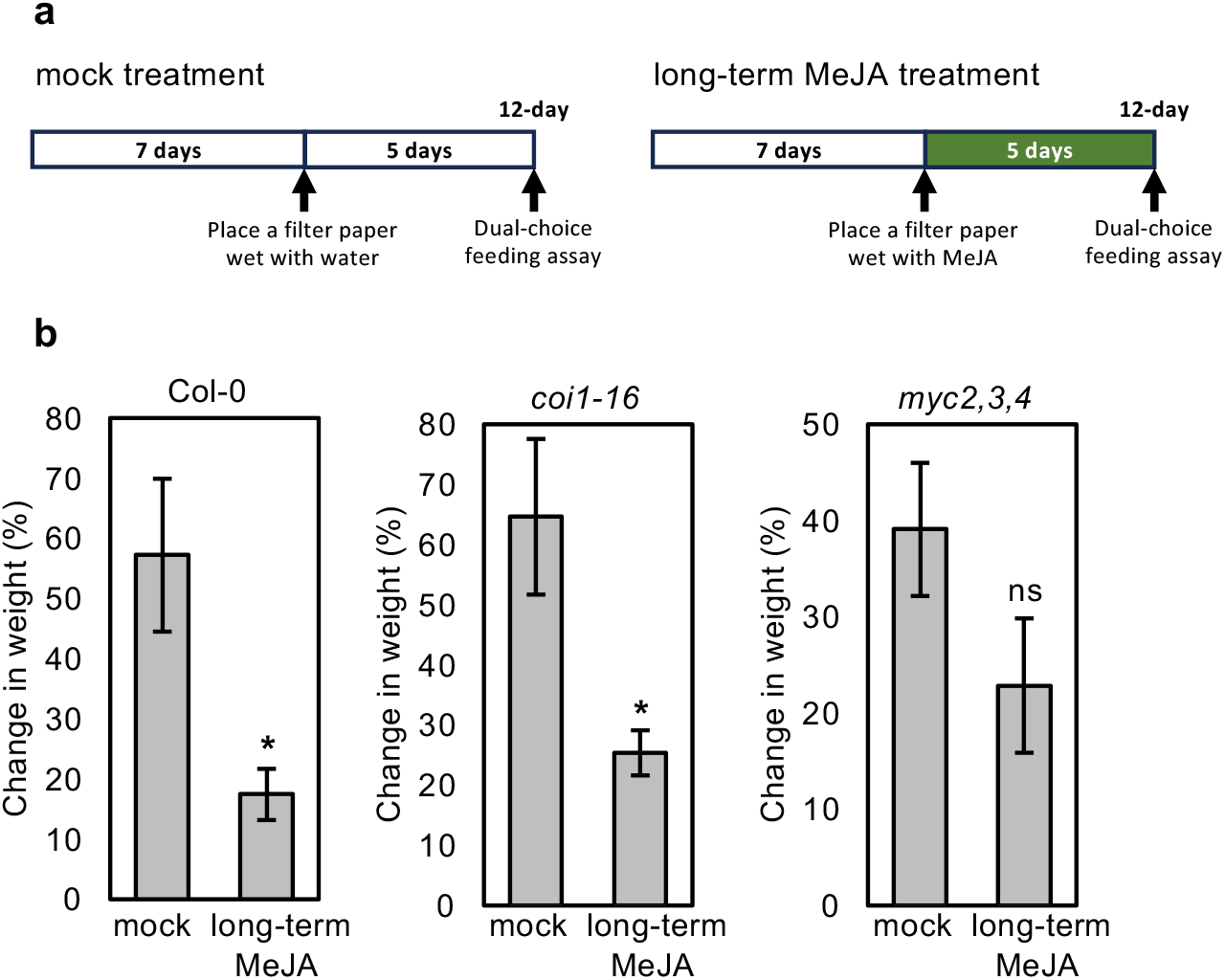
Effect of long-term MeJA treatment on cricket feeding preference. **a)** The schemes show the experimental setup of mock and long-term MeJA treatments. MeJA treatment was started 7 d after germination, and 12-d-old plants were subjected to the dual-choice feeding assay. **b)** The chart shows the ratio of plant weight reduction by feeding. Crickets of the same age, originating from the same colony, were starved for 2 d before the experiments. The 7-d-old Arabidopsis wild type plants, *coi1-16* and *myc2,3,4* mutants, were treated with either mock or MeJA for 5 d. Afterward, ten MeJA-treated and untreated plants were carefully transferred from their growing medium into a 20×20×20 cm box (a dual-choice feeding assay). Ten starved crickets were released into each box and kept overnight. Plant weights were recorded before and after the feeding experiments and calculated as a change in weight (%). Error bars indicate standard error of four independent experiments (*n* = 4). Significance values were calculated by a two-sided Student’s *t*-test. Asterisk denotes *p* < 0.05, and ns denotes no significance based on the Student’s *t*-test.

## 4 Discussion

In nature, plants are constantly subjected to stresses such as wounding or herbivore attacks, which induce JA biosynthesis and subsequently activate the JA signaling pathway (McConn et al., 1997; Halitschke and Baldwin, 2004). Exposure of plants to exogenous MeJA elicits stress responses by activating JA-dependent signaling pathways (Jiang et al., 2017). In this study, we found that airborne MeJA signals induce defense-related myrosinase genes, namely *TGG1* and *TGG2*, in *Arabidopsis* plants. However, we unexpectedly found that long-term MeJA treatment induced the expression of *TGG1* and *TGG2* independent of the canonical JA signaling pathway using COI1 and MYC2/3/4.

It has been previously reported that a 6 h MeJA treatment induces the expression of *TGG1* but reduces the expression of *TGG2* (Feng et al., 2021). Under our experimental conditions, we did not observe any significant changes in the expression of *TGG1* until 2 d of MeJA treatment. This discrepancy may be due to the method of MeJA treatment; in this study slowly evaporating MeJA was applied to the plants through the air inside the plates, whereas MeJA was directly applied to the plants in the study by Feng et al. (2021). In our time course experiment, the upregulation of *TGG1* and *TGG2* in wild type started after 2 d of MeJA treatment, whereas the increase of *TGG1* and *TGG2* in *coi1-16* started after 4 d and 2 d of MeJA treatment, respectively. These findings suggest that the COI1-independent pathway is activated after 2 d of MeJA treatment; however, the response of *TGG1* to the COI1-independent pathway is slower than that of *TGG2* and requires more time.

Long-term airborne MeJA treatment induces *TGG1* and *TGG2*, but not through COI1 and MYC2/3/4 signaling pathways. Under long-term MeJA treatment, *TGG1* and *TGG2* gene expression increased in both *coi1-16* and *myc2,3,4* mutants; however, typical JA responses, such as *VSP2* and *BGLU18* gene expression (Liu & Timko, 2021; Schweizer et al., 2013; Stefanik et al., 2020) and the induction of growth inhibition (Zhang & Turner, 2008), were attenuated in the mutants compared to the wild type. Therefore, the regulatory systems of *TGG1* and *TGG2* in response to long-term MeJA treatment are unique.

There is an example of COI1-independent JA responses in plants. JA inhibits the formation of lateral roots through the auxin action, and its inhibitory effect is still observed in *Arabidopsis coi1* mutant, indicating that the effect is COI1-independent (Ishimaru et al., 2018). In this phenomenon, JA prevents the degradation of AUXIN (Aux)/IAA repressor proteins. This JA activity on lateral root inhibition was reduced in *auxin signaling f-box protein 5* (*afb5*) mutant, suggesting that the AFB5 protein is involved in stabilizing Aux/IAA proteins in the COI1-independent JA signaling pathway (Ishimaru et al., 2018). Besides, the bypathing of the canonical hormone signaling pathway has been reported for auxin. F-box proteins TRANSPORT INHIBITOR RESPONSE1 (TIR1) form a co-receptor complex with Aux/IAA proteins, which is mechanically analogous to the COI1-JAZ co-receptor complex, as both plant hormone receptors function as ligand-induced degradation of repressor proteins. During the differential growth of the apical hook in the *Arabidopsis* seedlings, auxin promotes the cleavage of TRANSMEMBRANE KINASE 1 (TMK1) to release its kinase domain for the activation of downstream substrates, and this cleavage has been shown to be TIR1-independent (Cao, et al., 2019). TMK1 is also involved in other hormone signaling networks, such as abscisic acid (ABA); High concentrations of auxin enhanced ABA responses in a TMK1-dependent manner (Yang et al., 2021). These findings suggest the role of the TMK family proteins in alternative hormone signaling pathways, including auxin, ABA, and potentially other plant hormone signaling pathways.

MYC2/3/4 are TFs involved in JA signaling; however, our analysis indicated that MYC2/3/4 are not involved in the regulation of *TGG1* and *TGG2* expression in response to long-term MeJA treatment. FAMA is a key TF for myrosin and stomatal guard cell differentiation, and *TGG1* and *TGG2* expression (Li and Sack, 2014; Shirakawa et al., 2014; Feng et al., 2021). We observed increased *FAMA* mRNA levels in the first, second, and fourth rosette leaves, suggesting that FAMA is the TF for the expression of *TGG1* and *TGG2* after long-term MeJA treatment in our experiment. Due to the pleiotropic growth defect in *fama* mutants (Ohashi-Ito and Bergmann, 2006), we could not assess their response to long-term JA treatment. Nevertheless, our findings are consistent with previous observations that an increase in *FAMA* transcripts correlates with the activation of *TGG1* and *TGG2* (Feng et al., 2021), indicating that *FAMA* mRNA accumulation is a hallmark of the activation of downstream *TGG1* and *TGG2* gene expression. Our microscopic observation demonstrated consistent *FAMA* expression in stomatal guard cells. Long-term MeJA treatment increases *FAMA* expression in stomatal cells but not in myrosin cells in old leaves, suggesting that the increased expression of *TGGs* in old leaves comes from elevated expression of those genes in stomatal cells. Interestingly, in 12-d-old plants, myrosin cell expression of *FAMA* was not observed in the first and second leaves but was in the fourth leaf. Therefore, we propose a model in which the JA response differs depending on leaf age or phyllotaxy, with younger (late-emerged) leaves responding to JA in both stomatal guard cells and myrosin cells and older (early-emerged) leaves responding only to JA in stomatal guard cells (Supplementary Fig. 12).

We found a positive impact of long-term MeJA treatment on the myrosin cell area in young leaves at certain plant ages, indicating that this phenomenon positively correlates with the upregulation of *TGG1* and *TGG2* expression in shoots. Myrosin cell area expansion can be explained by two different mechanisms: one is by promoting myrosin cell differentiation to increase these cell numbers and the other is by promoting myrosin cell enlargement. MeJA treatment did not enhance myrosin cell numbers in young leaves (Supplementary Fig. 4), suggesting that the increase in myrosin cell area was not a consequence of differentiation of myrosin cells. Therefore, we speculate that long-term exposure to MeJA enhances myrosin cell expansion, along with *FAMA, TGG1*, and *TGG2* expressions at certain developmental stages or leaf orders. Supporting this hypothesis, Noir et al. (2013) reported that MeJA inhibits the mitotic cycle in the G1 phase before the S-phase transition in leaf and root cells, resulting in an increase in cell size.

In conclusion, this study highlights the complexity of MeJA signaling in plant defense. The expression patterns of *TGG1* and *TGG2* in *coi1-16* and *myc2,3,4* mutants suggest that multiple gene regulatory pathways are involved and that long-term MeJA treatment induces an additional pathway. Our analysis highlights the importance of the MeJA treatment method and duration of treatment in gene expression changes. Our study provides a new understanding underlying the regulation of myrosinases in response to MeJA.

## Author Contributions

M.M. and K.Y. designed the study. M.M. and A.P. performed the experiments. M.M. and S.G-Y. performed feeding experiments. M.M. and K.Y. analyzed the data. M.M., K.E., E.D., and K.Y. wrote the manuscript.

## Acknowledgements

We thank Makoto Shirakawa (Nara Institute of Science and Technology) and Ikuko Hara-Nishimura (Konan University) for providing the *ProTGG2:VENUS-2sc* and *pFAMA:GFP* transgenic lines and anti-TGG1 and anti-TGG2 antibodies, and Henning Frerigman (Max Plank Institute for Plant Breeding Research) for providing *myc2,3,4*. We are grateful to M. Shirakawa for fruitful discussions and comments on this study.

## Data availability statement

The data that support the findings of this study are available from the corresponding author upon request.

