## Supplementary Figures for "Long-term Exposure to Methyl Jasmonate Increases Myrosinases TGG1 and TGG2 in Arabidopsis *coi1* and *myc2,3,4* Mutants"

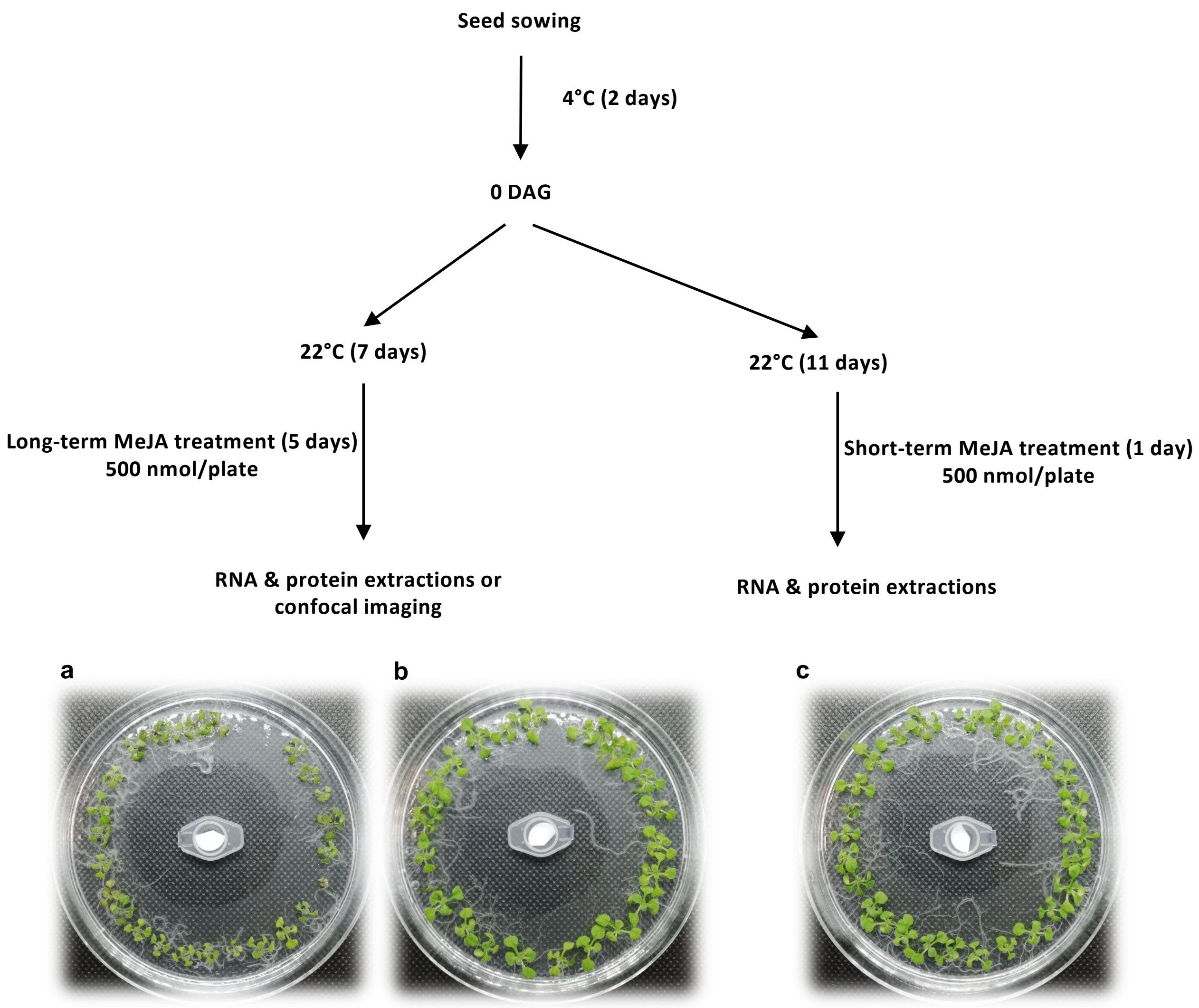


**Supplementary Figure 1. Airborne MeJA treatment method**.
The liquid of MeJA was poured onto a small paper pad in the cap of Eppendorf tube and positioned at the center of a plate ($\varphi$ = 9 cm), and plants were grown 4 cm away from the MeJA source (a, c). Distilled water was used instead of MeJA for the mock treatment (b). Wild-type plants treated with MeJA for 5 d (a) have smaller sizes compared to the mock-treated plants (b).

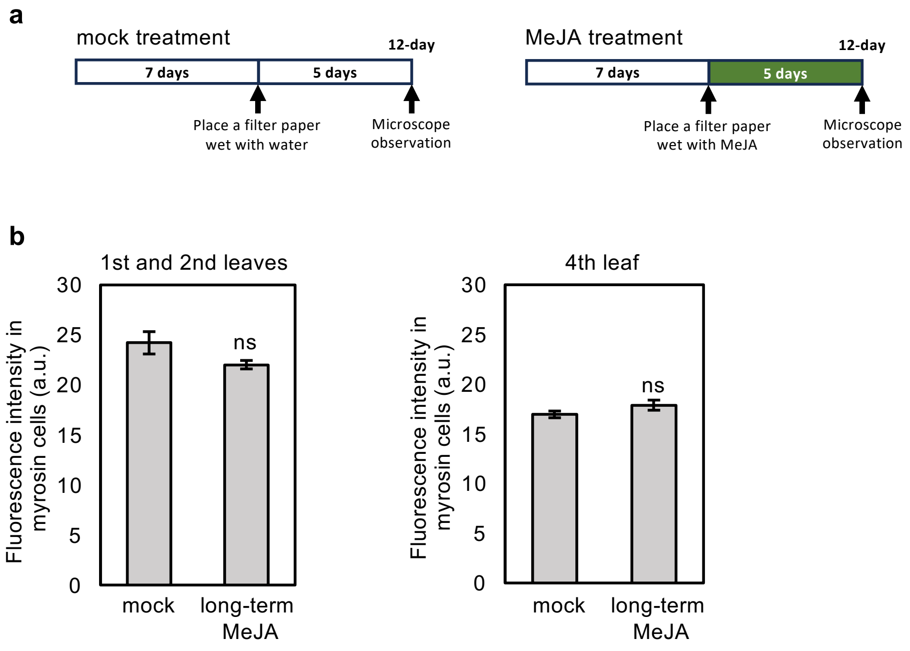


**Supplementary Figure 2. Long-term MeJA treatment does not change Venus fluorescence in myrosin cells.**

**a)** The schemes show the experimental setup of mock and long-term MeJA treatment. The treatment was conducted in the same way as in Figure 1a; MeJA treatment was started 7 d after germination, and plants were examined 12 d after germination. **b)** The fluorescent intensity in myrosin cells was calculated from the confocal microscope images of *pTGG2:Venus-2sc* transgenic plants. Error bars denote the standard error of seven biological replications. ns denotes no significance based on the Student's *t*-test.

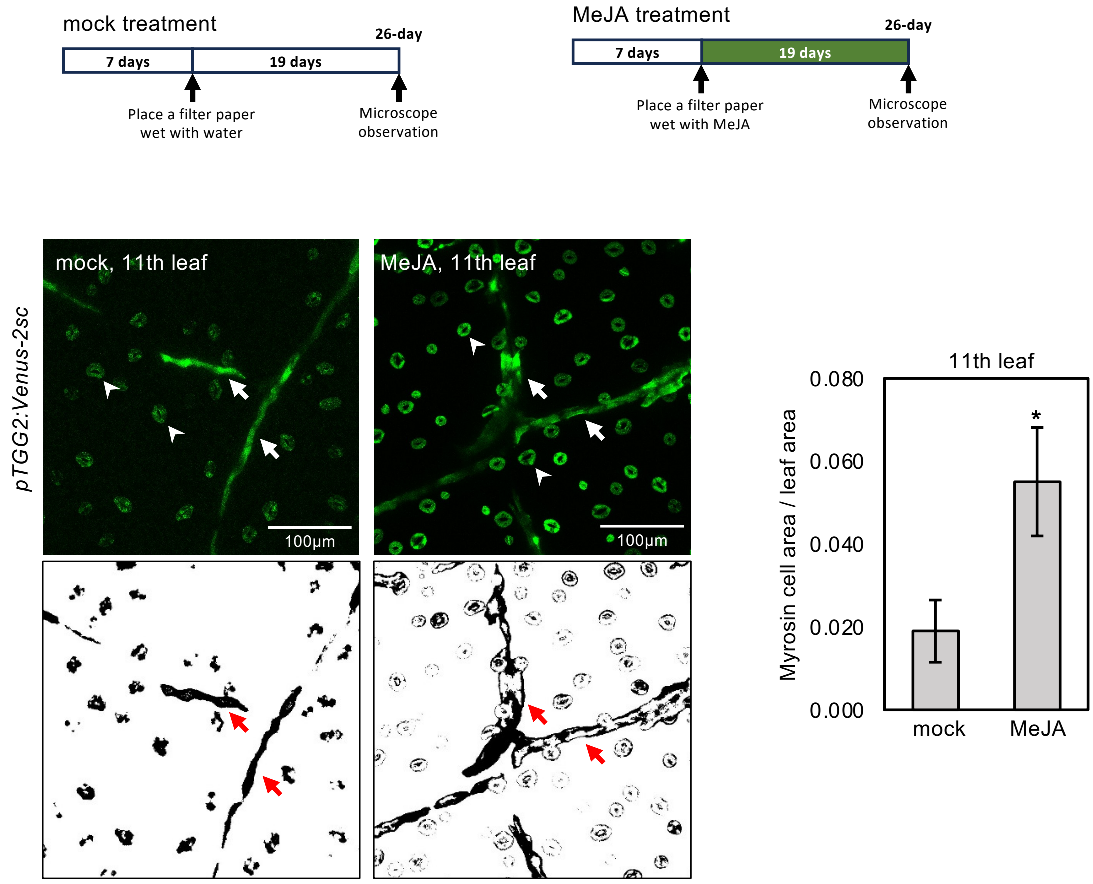


**Supplementary Figure 3. Long-term MeJA treatment enhances myrosin cell area in younger leaves.**

Confocal microscopic images of the eleventh leaves of 12-d-old transgenic plants harboring *pTGG2:Venus-2sc*. The chart shows the ratio of myrosin cell areas per leaf area in the eleventh leaves. The MeJA treatment was prolonged for 19 d. Error bars denote the standard error of seven biological replications. * denotes $p < 0.05$ based on the Student's *t*-test.

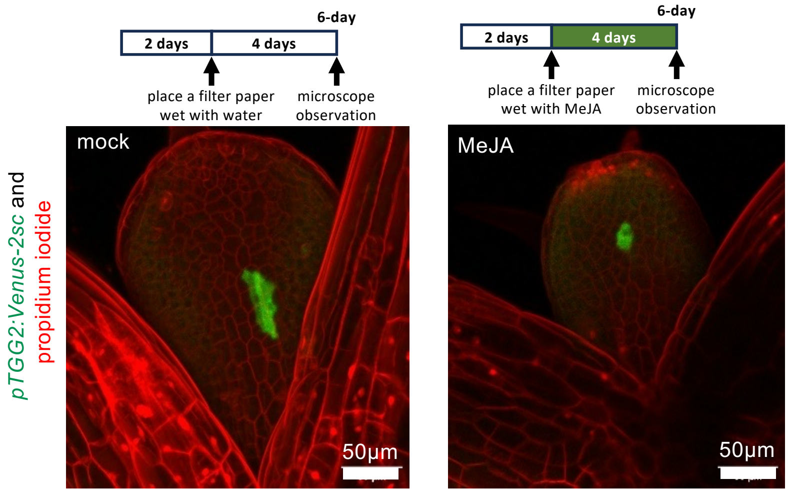


**Supplementary Figure 4. MeJA treatment does not change number of myrosin cells in emerging first leaves.**

Confocal microscopic images of the emerging first leaves of 6-d-old transgenic plants harboring pTGG2:Venus-2sc. MeJA treatment was started 2 d after germination. Green shows Venus fluorescence in myrosin cells, and red shows propidium iodide fluorescence in cell walls and nuclei.

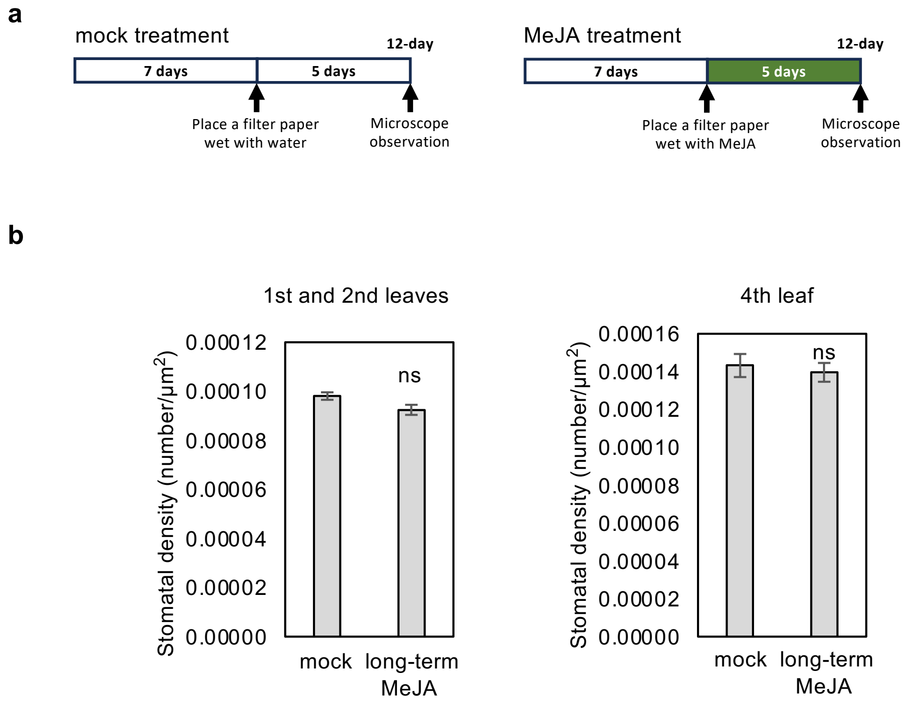


**Supplementary Figure 5. Long-term MeJA treatment does not change stomatal density.**

**a)** The schemes show the experimental setup of mock and long-term MeJA treatment. The treatment was conducted in the same way as in Figure 1a; MeJA treatment was started 7 d after germination, and plants were examined 12 d after germination. **b)** Stomatal density was calculated from the confocal microscope images of *pTGG2:Venus-2sc* transgenic plants. Error bars denote the standard error of five biological replications. ns denotes no significance based on the Student's *t*-test.

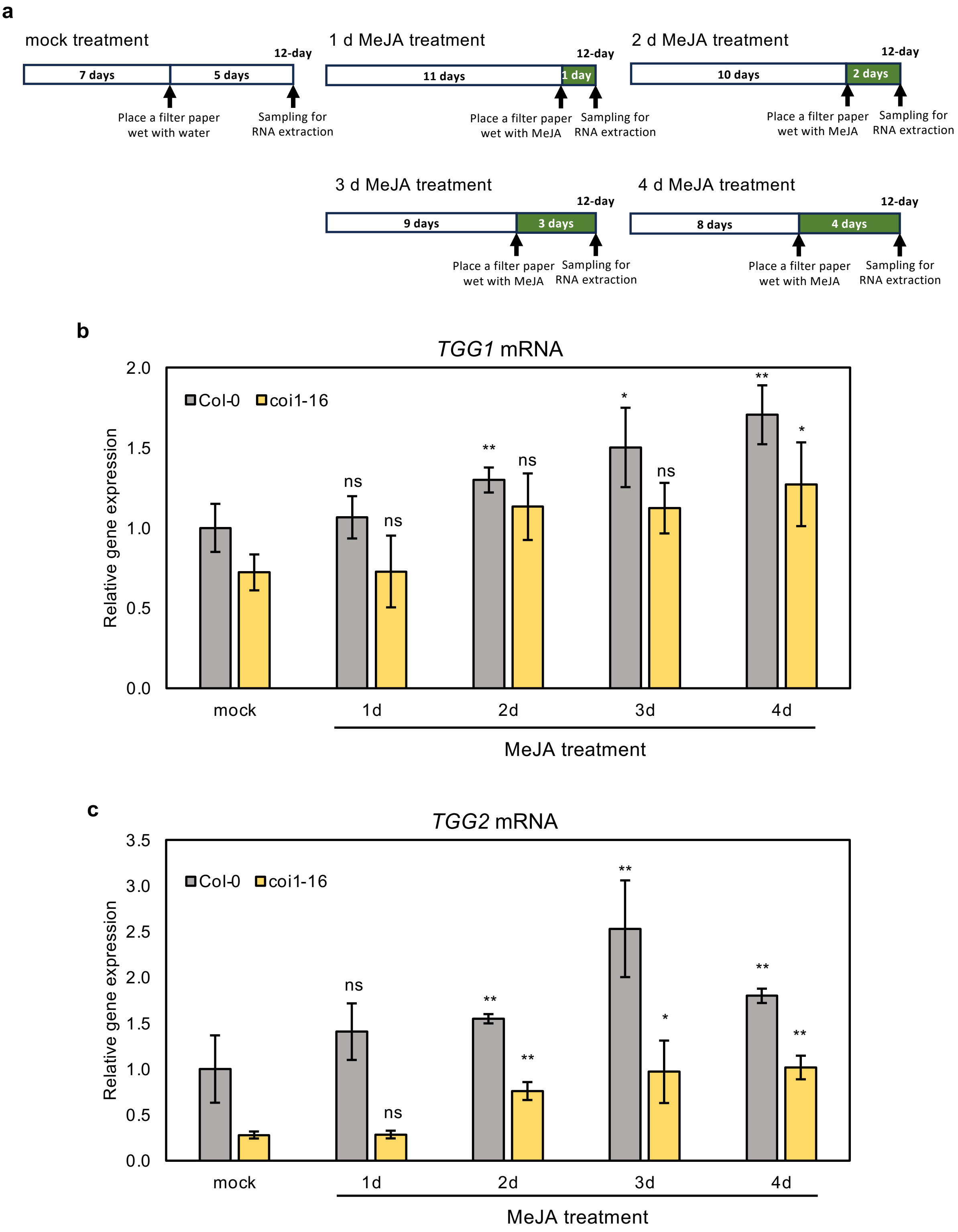


**Supplementary Figure 6. Transcript level of *TGG1* and *TGG2* in Col-0 and *coi1-16* mutant.**
**a)** The schemes show the experimental setup of mock and MeJA treatments. **b, c)** The relative expression levels of *TGG1* (b) and *TGG2* (c) in leaves of mock and 1 d, 2 d, 3 d, and 4 d after MeJA treatment in Col-0 and *coi1-16* mutant. Error bars denote the standard error of three biological replications. ns denotes no significance, * denotes $p < 0.05$ and ** denotes p < 0.01 based on the Student's *t*-test.

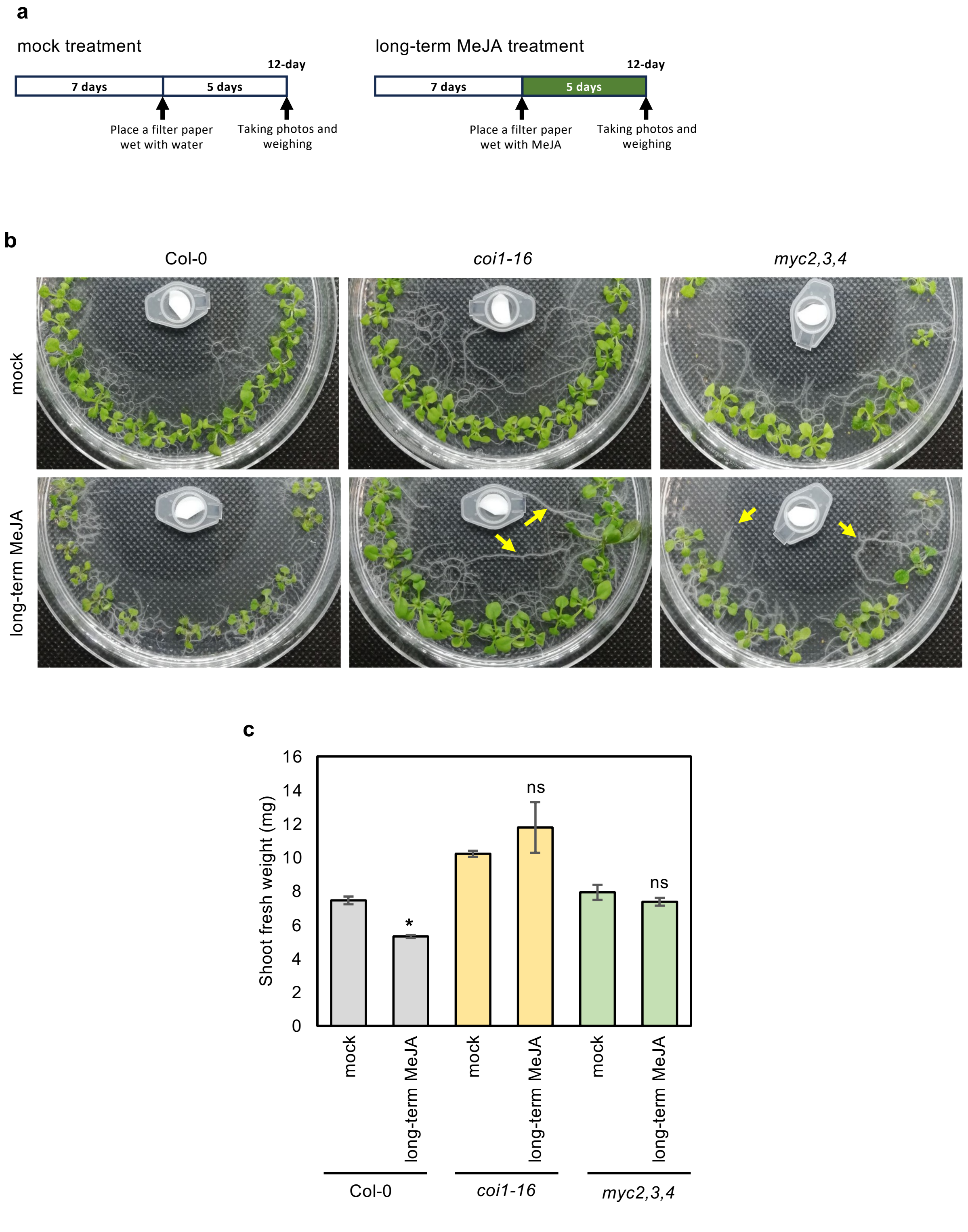


**Supplementary Figure 7. MeJA treatment does not reduce plant growth in *coi1-16* and *myc2,3,4* mutants.** **a)** The schemes show the experimental setup of mock and MeJA treatments. **b)** Images of mock or long-term MeJA treated 12-d-old-plants. Arrows show root growth. **c)** Shoot fresh weight of plants used in (b). Error bars denote the standard error of ten plants. Asterisk denotes $p < 0.05$, and ns denotes no significance based on the student's *t*-test.

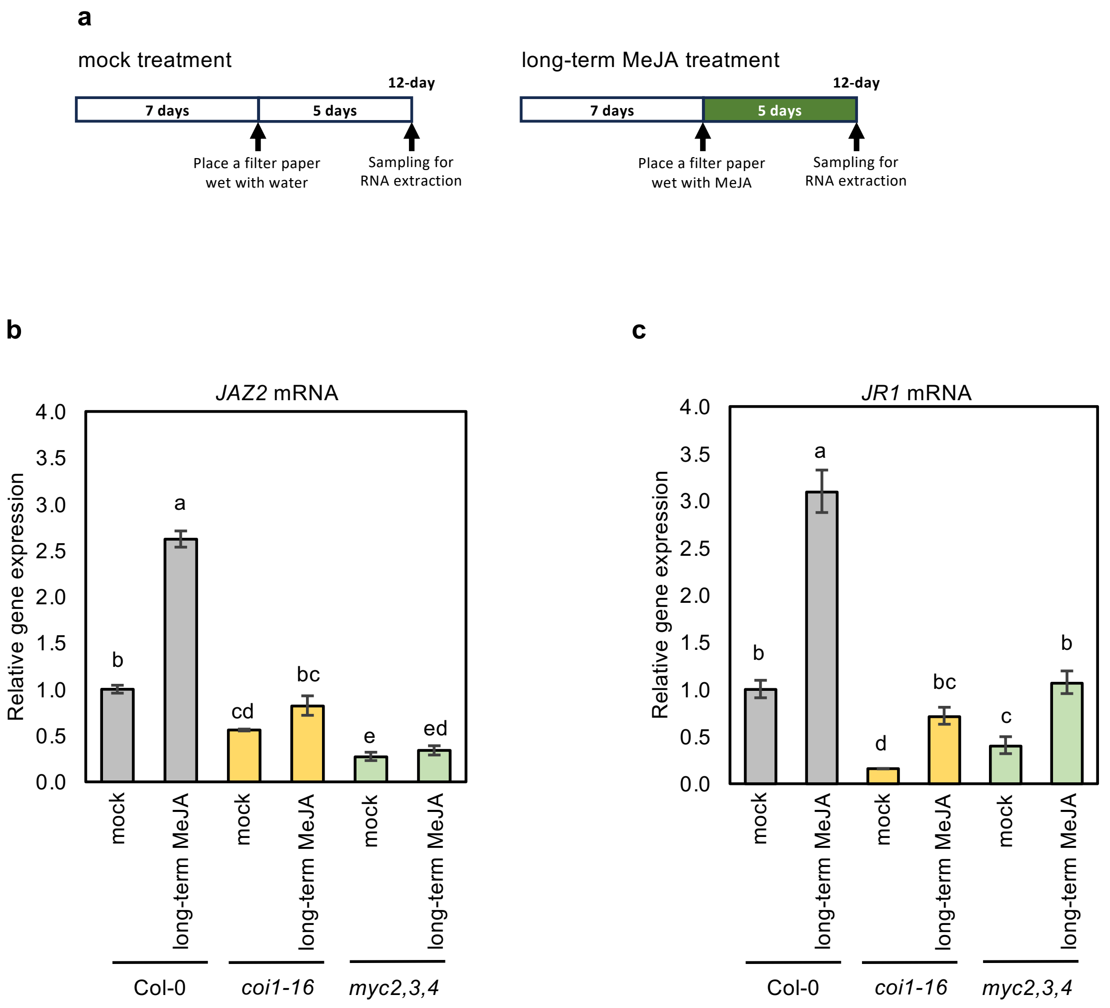


**Supplementary Figure 8. Expression of Canonical MeJA-responsive genes decreased significantly in *coi1-16* and *myc2,3,4* mutants.**

**a)** The schemes show the experimental setup of mock, short-term and long-term MeJA treatments. MeJA treatment was started 7 d after germination, and plants were sampled 12 d after germination. **b, c)** The relative expression levels of *JAZ1* (b), and *JR1* (c) in the shoots of mock and long-term MeJA-treated Arabidopsis wild type (Col-0), *coi1-16* and *myc2,3,4* mutants. Error bars indicate SE (n = 3 replicates). Different lowercase letters indicate significant differences ($p < 0.05$; Tukey's test).

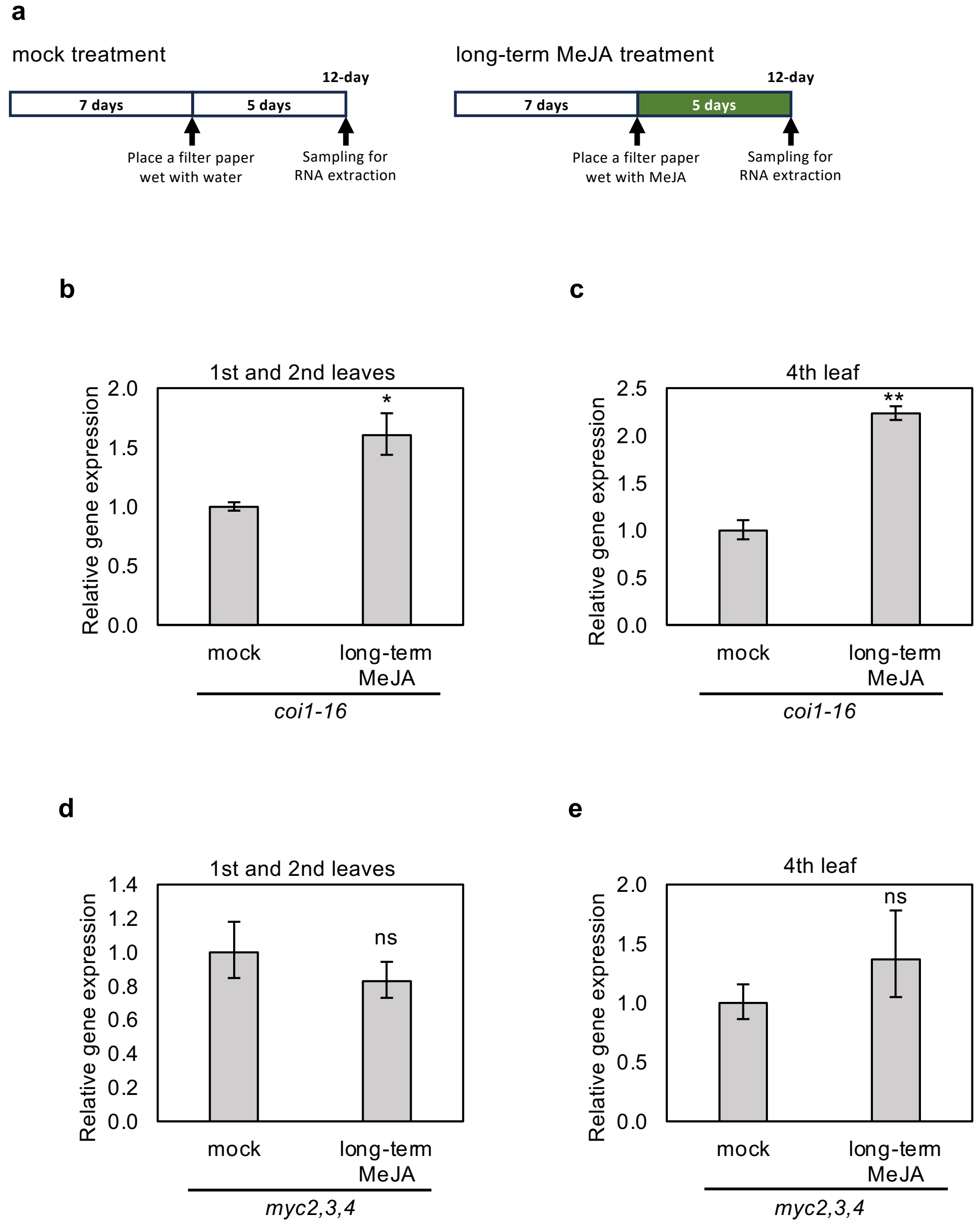


**Supplementary Figure 9. Long-term MeJA treatment increased the expression of *FAMA* in the first, second and fourth leaves of *coi1-16* mutant, while no significant change was observed in *myc2,3,4* mutant.**

**a)** The schemes show the experimental setup of mock, short-term and long-term MeJA treatments. MeJA treatment was started 7 d after germination, and plants were sampled 12 d after germination. **b, c)** The relative expression levels of *FAMA* in different rosette leaves of 12-d-old *coi1-16* mutant. **d, e)** The relative expression levels of *FAMA* in different rosette leaves of 12-d-old *myc2,3,4* mutant. Error bars denote the standard error of three biological replications. ns denotes no significance, * and ** denotes $p < 0.05$ and $p < 0.01$ based on the student's $t$-test.

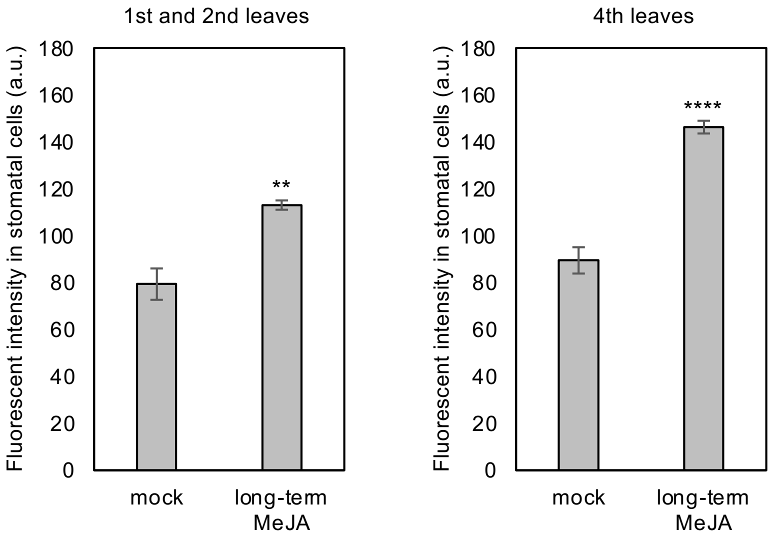


**Supplementary Figure 10. Long-term MeJA treatment increased *FAMA* expression in the stomatal cells of first, second and fourth leaves.**

The GFP fluorescent intensity in stomatal cells was calculated from the confocal microscope images of *pFAMA:GFP* transgenic plants. Error bars denote the standard error of seven biological replications. ** and **** denote respectively $p < 0.01$ and $p < 0.0001$ based on the Student's *t*-test.

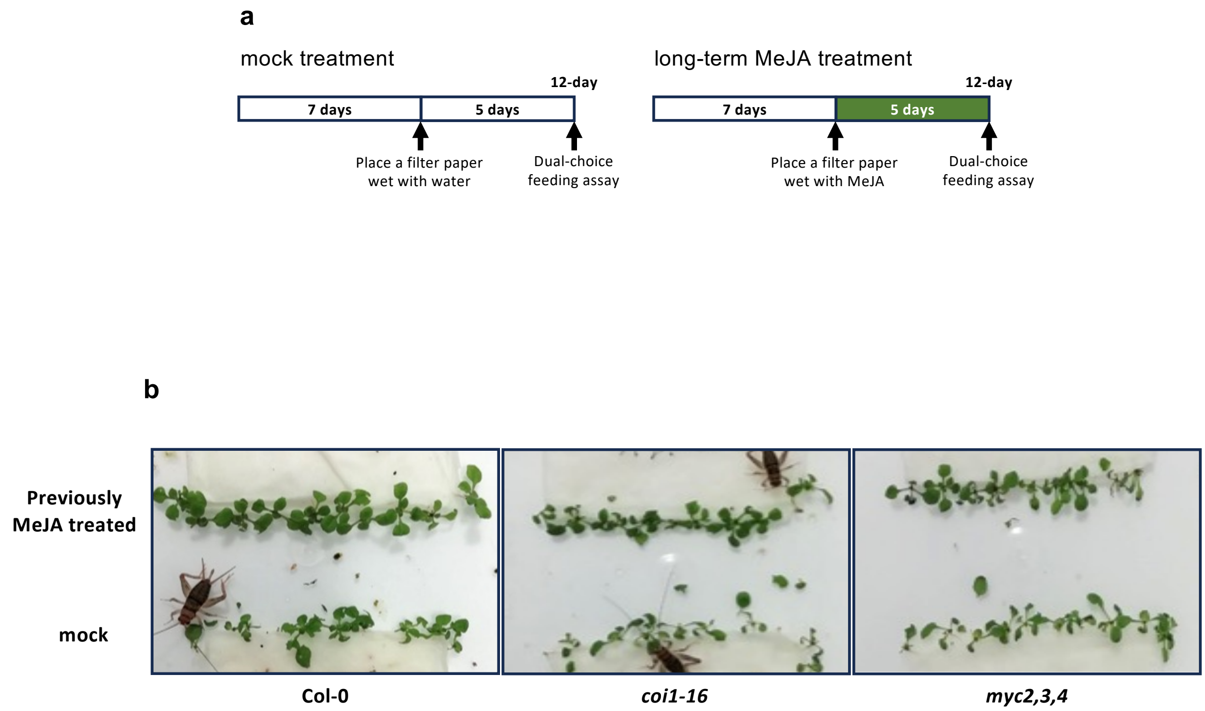


**Supplementary Figure 11. Set up of crickets feeding assay.**

**a)** The schemes show the experimental setup of mock and long-term MeJA treatments. MeJA treatment was started 7 d after germination, and 12-d-old plants were subjected to the dual-choice feeding assay. **b)** The photos show our setup for the dual choice cricket feeding assay in wild type, *coi1-16*, and *myc2,3,4* mutants. Ten MeJA-treated and untreated plants were carefully transferred from their growing medium into a 20×20×20 cm box (a dual-choice feeding assay). Ten starved crickets were released into each box. The images show representatives of four individual experiments in each plant line.

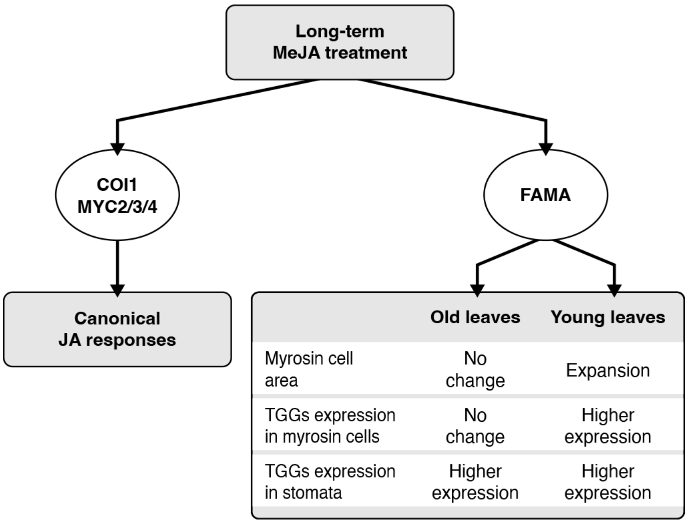


**Supplementary Figure 12. Schematic model for the effects of long-term MeJA treatment on myrosinases in old and young leaves.**

**Supplementary Table 1. List of primers used in this study**

| Name | Sequence (5' to 3') |
|---|---|
| qPCR_TGG1-FP | GCAGCGGCCGTTGATGTTTA |
| qPCR_TGG1-RP | TAGCCCGCTCAGTTGCATCT |
| qPCR_TGG2-FP | TGGAGCCGCTAACAAAGGGT |
| qPCR_TGG2-RP | AGGCGAGCTTCTGTGCTGTT |
| qPCR_FAMA-FP | ACTACGGTAGCGAACCAAGC |
| qPCR_FAMA-RP | CGACTTGTTCTCCGCAGTCT |
| qPCR_BGLU18-FP | TGAGTGGCAAGATGGGTACA |
| qPCR_BGLU18-RP | TCAGCTTGGAGGTTGGAAAC |
| qPCR_VSP2-FP | CGTCGATTCGAAAACCATCT |
| qPCR_VSP2-RP | GGCACCGTGTCGAAGTCTAT |
| qPCR_JR1-FP | TCCTTCGTCCACACCATTGA |
| qPCR_JR1-RP | CTCCACCATCTCCACCTTGT |
| qPCR_JAZ2-FP | CTCTTTAGCCTGCGAACTCC |
| qPCR_JAZ2-RP | TTGGTATGGTGCCTTTGATG |
| UBQ10-Q2-FP | GAAGTGGAAAGCTCCGACAC |
| UBQ10-Q2-RP | TTAGAAACCACCACGAAGACG |
